# Metabolic Flux Hierarchy Prioritizes the Entner-Doudoroff Pathway for Carbohydrate Co-Utilization in *Pseudomonas protegens* Pf-5

**DOI:** 10.1101/402073

**Authors:** Rebecca A. Wilkes, Caroll M. Mendonca, Ludmilla Aristilde

## Abstract

The genetic characterization of *Pseudomonas protegens* Pf-5 was recently completed. However, the inferred metabolic network structure has not yet been evaluated experimentally. Here we employed ^13^C-tracers and quantitative flux analysis to investigate the intracellular network for carbohydrate metabolism. Similar to other *Pseudomonas* species, *P. protegens* Pf-5 relied primarily on the Entner-Doudoroff (ED) pathway to connect initial glucose catabolism to downstream metabolic pathways. Flux quantitation determined that, in lieu of the direct phosphorylation of glucose by glucose kinase, phosphorylation of oxidized products of glucose (gluconate and 2-ketogluconate) towards the ED pathway accounted for over 90% of consumed glucose and greater than 35% of consumed glucose was secreted as gluconate and 2-ketogluconate. Consistent with the lack of annotated pathways for the initial catabolism of pentoses and galactose in *P. protegens* Pf-5, only glucose was assimilated into intracellular metabolites in the presence of xylose, arabinose, or galactose. However, when glucose was fed simultaneously with fructose or mannose, co-uptake of the hexoses was evident but glucose was preferred over fructose (3 to 1) and over mannose (4 to 1). Despite gene annotation of mannose catabolism toward fructose 6-phosphate, metabolite labeling patterns revealed that mannose-derived carbons specifically entered central carbon metabolism via fructose-1,6-bisphosphate, similarly to fructose catabolism. Remarkably, carbons from mannose and fructose were found to cycle backward through the upper Emden-Meyerhof-Parnas pathway to feed into the ED pathway. Therefore, the operational metabolic network for processing carbohydrates in *P. protegens* Pf-5 prioritizes flux through the ED pathway to channel carbons to downstream metabolic pathways.

**IMPORTANCE:** Species of the *Pseudomonas* genus thrive in various nutritional environments and have strong biocatalytic potential due to their diverse metabolic capabilities. Carbohydrate substrates are ubiquitous both in environmental matrices and in feedstocks for engineered bioconversion. Here we investigated the metabolic network for carbohydrate metabolism in *P. protegens* Pf-5. Metabolic flux quantitation revealed the relative involvement of different catabolic routes in channeling carbohydrate carbons through the network. We also uncovered that mannose catabolism was similar to fructose catabolism, despite the gene annotation of two different pathways in the genome. Elucidation of the constitutive metabolic network in *P. protegens* is important for understanding its innate carbohydrate processing, thus laying the foundation for targeting metabolic engineering of this untapped *Pseudomonas* species.

## INTRODUCTION

Species of the genus *Pseudomonas,* which are ubiquitous in the environment, are metabolically diverse and often used for industrial bioproduction (1). Elucidating the native network of carbon fluxes through metabolic pathways is critical to the engineering of these bacterial species to optimize their use in agriculture, industry, and medicine. Gaining importance in bioremediation, *P. protegens* Pf-5 was identified to produce enzymes that degrade polyurethane, a plastic polymer (2). Furthermore, *P. protegens* Pf-5 is also known to synthesize and release several antimicrobials and exoenzymes that are toxic to plant pathogens (3–6). Recently, *P. protegens* Pf-5 was characterized and annotated at the genomic level (7). However, the metabolic network of *P. protegens* Pf-5 has only been inferred from genome annotation and has not yet been investigated experimentally.

Given the importance and ubiquity of carbohydrate-containing feedstocks, we seek to unravel the metabolic network structure for carbohydrate metabolism in *P. protegens* Pf-5 by combining ^13^C-assisted cellular carbon mapping with ^13^C metabolic flux analysis (MFA). Previous studies on other *Pseudomonas* species (i.e. *P. putida, P. fluorescens*) have focused on elucidating metabolic fluxes during growth on glucose, a prototypical carbohydrate substrate (8–11). In a similar fashion, we also studied here the innate carbohydrate metabolism in *P. protegens* Pf-5 during feeding on glucose alone. However, carbon feedstocks are typically composed of other carbohydrates in addition to glucose. Therefore, we also investigated carbon assimilation and fluxes when the *P. protegens* Pf-5 cells were fed on mixtures of glucose with other hexoses (mannose, fructose, and galactose) or pentoses (xylose and arabinose).

Previous reports showed that *P. protegens* strains were able to grow on glucose, mannose, or fructose as a single carbon source, but not on galactose, xylose, or arabinose (6). In the genome of *P. protegens* Pf-5, the genes for the following transporters were encoded: a phosphoenolpyruvate (PEP):sugar phosphotransferase system (PTS) for fructose uptake (and possibly for mannose uptake) and ATP-binding cassette (ABC) transporters for glucose, mannose, galactose, and xylose (Fig. 1; Table S1) (7). Characteristically in *Pseudomonas* species, glucose metabolism involves a peripheral pathway in the periplasm wherein glucose dehydrogenase converts glucose to gluconate and gluconate 2-dehydrogenase converts gluconate to 2-ketogluconate (Fig. 1) (8). Fluxes through these oxidation reactions were found to maximize growth on glucose (8). After active transport into the cytoplasm, the oxidized products of glucose are phosphorylated to 6-phosphogluconate (6P-gluconate) and subsequently routed to the Entner-Doudoroff (ED) pathway or the pentose phosphate (PP) pathway (Fig. 1) (8, 10, 11). Previous studies with other *Pseudomonas* species have also determined that the forward Embden-Meyerhof-Parnas (EMP) pathway is not possible due to the absence of 6-phosphofructokinase to convert fructose 6-phosphate (F6P) to fructose 1,6-bisphosphate (FBP) (Fig. 1) (1, 7, 10). Therefore, to route glucose-derived carbons eventually downstream towards biosynthetic pathways and the tricarboxylic acid (TCA) cycle, the ED pathway is required wherein 6P-gluconate is cleaved to produce pyruvate and glyceraldehyde 3-phosphate (GAP) (Fig. 1). The gene that encodes 6-phosphofructokinase is also absent in *P. protegens* Pf-5 (7), thus the ED pathway is assumed to be the required route for glucose metabolism in *P. protegens* Pf-5. However, the extent to which the peripheral pathway of glucose oxidation contributes to initial glucose catabolism compared to direct glucose phosphorylation in *P. protegens* Pf-5 remains to be determined.

**FIG 1.**
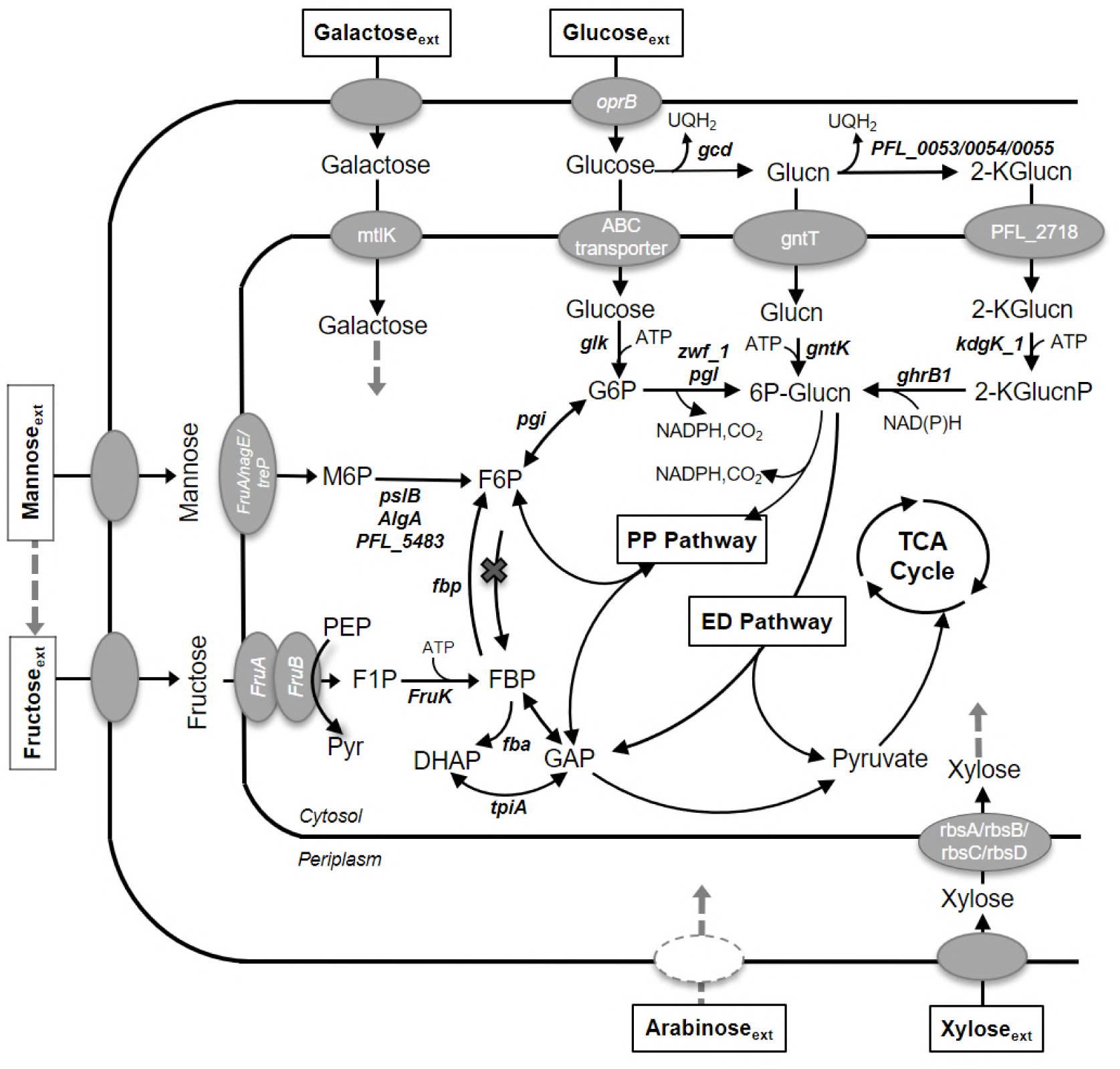
Putative genes involved in the uptake and initial catabolism of glucose, galactose, mannose, fructose, xylose, and arabinose into central metabolism. The gene annotations for pathways were collected from KEGG database (37–39) and MetaCyc database (40) for *P. protegens* Pf-5. The corresponding gene loci for the genes in the figure are shown in Table S1. The abbreviations are as follows: gluconate, Glucn; 2-ketogluconate, 2-KGlucn; 2-keto-6-phosphogluconate, 2-KGlucnP; glucose 6-phosphate, G6P; 6-phosphogluconate, 6P-Glucn; fructose 6-phosphate, F6P; fructose 1,6-bisphosphate, FBP; dihydroxyacetone-3-phosphate, DHAP; glyceraldehyde 3-phosphate, GAP; fructose 1-phosphate, F1P; pyruvate, Pyr; phosphoenolpyruvate, PEP.

In contrast to glucose, fructose is transported through the PTS, which uses the phosphate group from PEP to phosphorylate fructose to fructose-1-phosphate (F1P) followed by a subsequent phosphorylation step by 1-phosphofructokinase to convert F1P to FBP (Fig. 1) (9, 12, 13). Previous studies on other *Pseudomonas* species (*P. fluorescens, P. putida, P. aeruginosa, P. stutzeri,* and *P. acidovorans*) reported that fructose-derived carbons were cycled back via a reverse flux through upper EMP pathway (FBP up to G6P) to be connected to the ED pathway (12, 14–16). However, according to the genome of *P. protegens* Pf-5, FBP could also be subjected to the last step of the EMP pathway wherein FBP can be lysed directly to two triose-phosphates, GAP and dihydroxyacetone-3-phosphate (DHAP) (Fig. 1) (7). Whether the preferential route for fructose assimilation during simultaneous feeding on glucose is via the lower EMP pathway or through the reverse cycling of carbons from FBP to ED pathway remains to be determined. The catabolic routing of FBP has important energetic implications for *P. protegens* Pf-5. Compared to the direct lysis through the forward EMP pathway, the combination of reverse flux through upper EMP pathway with the ED pathway maintains the same quantity of reduced equivalents (i.e., NAD(P)H) but half the ATP yield.

With respect to initial mannose catabolism, a previous study with *P. aeruginosa* proposed two possible routes, which involve either mannose isomerization to fructose followed by subsequent phosphorylation to FBP or direct phosphorylation of mannose to mannose 6-phosphate (M6P) prior to isomerization to F6P (17, 18). Relevant to the first route, an intracellular mannose isomerase (EC 5.3.1.7) was reported in *P. cepacia* and *P. saccharophila* (19, 20), but the gene for this enzyme was not annotated in the *P. protegens* Pf-5 genome (7). On the other hand, albeit not yet confirmed by metabolic studies, the genes for the relevant enzymes in the second catabolic route, i.e. the conversion of mannose to F6P, were annotated in the *P. protegens* Pf-5 genome (Fig. 1).

The presence of the annotated genes for transketolase and transaldolase enzymes implied a fully functional PP pathway in *P. protegens* Pf-5 (7). Both oxidative and non-oxidative routes of the PP pathway are important to channel carbohydrate-derived metabolite precursors to the biosynthesis of ribonucleotides and aromatic amino acids. Regarding xylose catabolism, despite the annotation of a ribose transporter that could be used as a possible xylose transporter, the genes encoding the enzymes (xylose isomerase and xylulose kinase) responsible for introducing xylose into the PP pathway were not present in *P. protegens* Pf-5 (Fig. 1) (7). Moreover, the collective enzymes needed for the alternative route for xylose through the Weimberg pathway, which incorporates xylose through xylonate into α-ketoglutarate, were not all present in the genome of *P. protegens* Pf-5 (7). Regarding arabinose catabolism, there was no annotated pathway for the assimilation of arabinose in *P. protegens* Pf-5 (Fig. 1) (7). Despite the lack of the genes for xylose catabolism, a recent study reported extracellular xylose depletion by *P. protegens* Pf-5 during growth on a mixture of carbohydrates (21). Whether arabinose or xylose is incorporated into cellular metabolism, specifically the PP pathway, in the presence of another carbohydrate remains to be determined.

Here we applied liquid chromatography (LC) with high-resolution mass spectrometry (HRMS) to perform a ^13^C-assisted metabolomics investigation during growth of *P. protegens* Pf-5 on glucose alone or simultaneously with fructose, mannose, galactose, xylose, or arabinose. We provide the first quantitative evaluation of the hypothetical metabolic network of *P. protegens* Pf-5 deduced from its genome-level characterization. First, we employed carbon mapping to identify the specific pathways that channel glucose-derived carbons throughout cellular metabolism. Second, we performed quantitative analysis to determine energetic yields from the cellular metabolism in *P. protegens* Pf-5. Third, we determined which carbohydrates can be co-assimilated with glucose in *P. protegens* Pf-5 and subsequently quantified the metabolic fluxes when co-utilization occurred. Our findings provide new metabolic insights, which both resolve discrepant metabolic predictions from genome annotation and quantify fluxes in the metabolic network structure for carbohydrate processing in *P. protegens* Pf-5.

## RESULTS

### Physiological Parameters of *P. protegens* Pf-5 Grown on Carbohydrate Mixtures

We investigated the growth phenotype of *P. protegens* during feeding on different hexose combinations. Our growth rates were in close agreement with reported values for glucose-grown *P. putida* (0.56 ± 0.02 h^-1^) and *P. fluorescens* (0.49 ± 0.03 h^-1^) (8, 22). At a total carbon equivalence of 100 mM C, the growth rate of cells fed on glucose alone (0.56 ± 0.09 h^-1^) was similar to the growth rate of cells fed on a 1:1 mixture of glucose and fructose (0.52 ± 0.06 h^-1^) or glucose and mannose (0.50 ± 0.04 h^-1^) (Fig. 2A; Fig. S1; Table S2). We also found that the growth rate remained relatively unchanged when the cells were grown on 50 mM C glucose alone (0.47 ± 0.02 h^-1^) (Fig. 2A). Therefore, *P. protegens* would not be subjected to carbon limitation during growth on carbohydrate mixtures if either fructose or mannose was not assimilated from the mixture with glucose (Fig. 2A).

**FIG 2.**
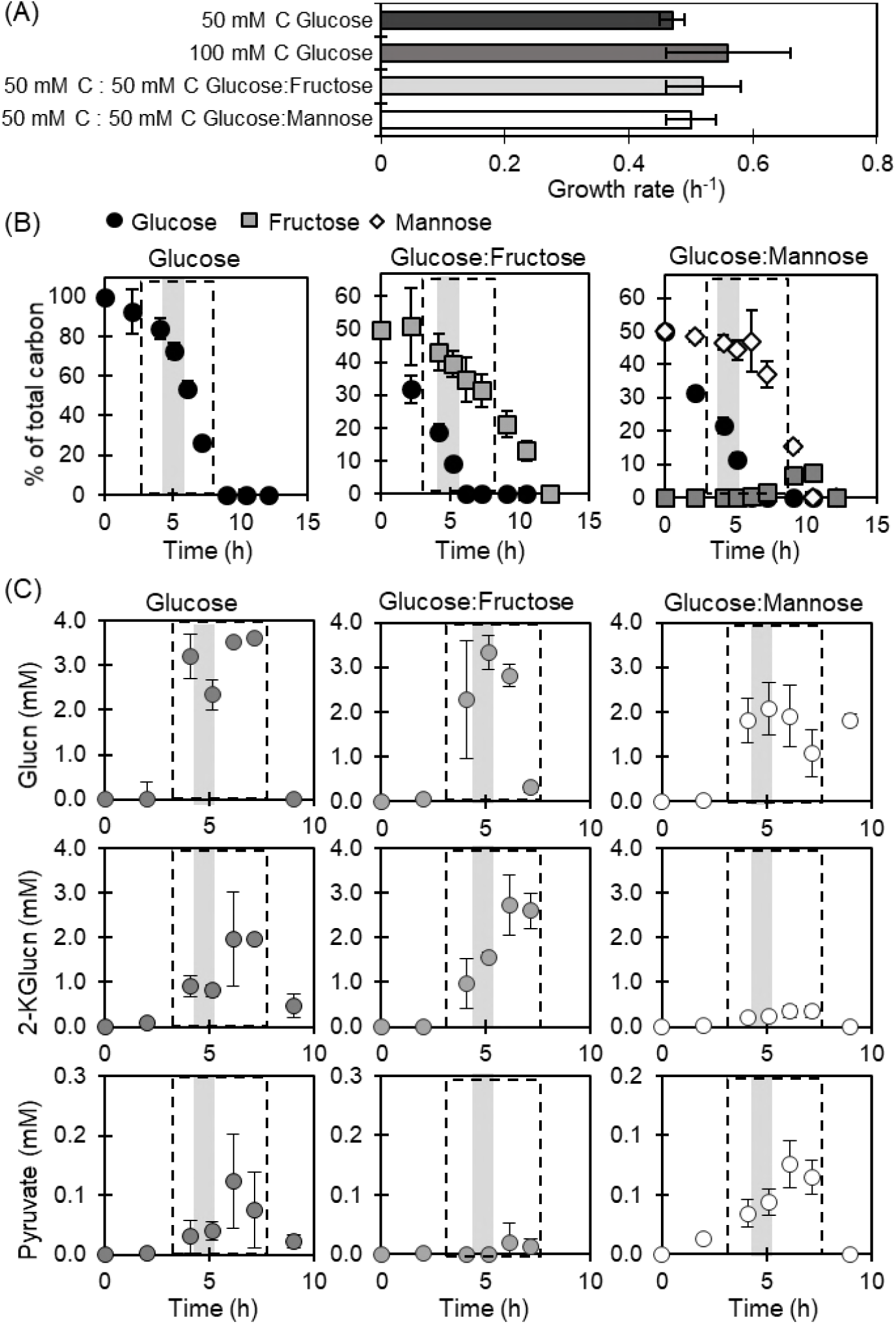
(A) Growth rate, (B) kinetic profile of carbohydrate depletion, and (C) kinetic profile of metabolite secretions for cells fed glucose only or equimolar glucose:fructose or glucose:mannose. Initial total carbohydrate concentration was 100 mM C (3 g L^-1^). In B and C, the data points within the dashed box were obtained during exponential growth; the shaded area at an OD_600_ of 0.5 corresponds to the timepoint for the measured labeling data used for the MFA. Data are shown as means ± standard deviation from biological replicates (*n* = 3).

We monitored substrate consumption by *P. protegens* Pf-5 cells by tracking the depletion of the carbohydrates from the extracellular medium (Fig. 2B; Table S2). The glucose-grown cells depleted glucose completely over a 10 h period (Fig. 2B). Over the same period of time, the cells grown on the mixture of glucose and mannose depleted both substrates completely but cells grown on the mixture of glucose and fructose depleted glucose completely and fructose partially (Fig. 2B). We also observed that, during growth of *P. protegens* Pf-5 on the glucose:mannose mixture, fructose appeared in the extracellular medium about 1 h after mannose consumption started (Fig. 2A). The appearance of extracellular fructose implied the presence of mannose isomerase, which was responsible for converting mannose to fructose in other *Pseudomonas* species (19,20) (Fig. 2B). During growth on both hexose mixtures, glucose was consumed faster and depleted by 6 h of growth, at which time about 30% of the fructose was consumed but only 10% of mannose was taken up by the cells (Fig 2B). Quantification of the consumption rates during exponential growth determined that fructose consisted 22% of the total carbon uptake and mannose consisted of 20% of total carbon uptake (Table S2). Thus, during hexose co-utilization under both of our experimental conditions, glucose acts as the major carbon source for cellular metabolism.

We also monitored the extracellular overflow of metabolic products, a phenomenon that is widely reported in *Pseudomonas* species (8, 11, 23). Both oxidized products of glucose (i.e., gluconate and 2-ketogluconate) and the organic acid pyruvate were found in appreciable levels (above 0.001 mM) in the extracellular medium (Fig. 2C). Secretions of gluconate and 2-ketogluconate were also reported with *P. putida* (8, 11) and *P. fluorescens* (23); pyruvate secretion was also reported in *P. fluorescens* (23). The levels of these secreted metabolites were dependent on substrate composition in the growth medium of *P. protegens*. The highest secretions of gluconate and 2-ketogluconate (greater than 2 mM) were measured in the medium when cells were grown on glucose alone or glucose with fructose. By contrast, during growth on the glucose:mannose mixture, the highest secretion of gluconate and 2-ketogluconate decreased substantially, by ∼40% and ∼85%, respectively (Fig. 2C; Table S2). Compared to gluconate and 2-ketogluconate, pyruvate was secreted in smaller amounts (μM range), with the highest secretion (0.13 ± 0.08 mM) obtained when cells were grown on glucose alone (Fig. 2C; Table S2).

### Metabolic Pathways and Fluxes in Glucose-Fed Cells

Isotopic enrichment with [1,2-^13^C_2_]-glucose was used to determine the metabolic network structure through initial glucose catabolism, the EMP pathway, the ED pathway, the PP pathway, and the TCA cycle (Fig. 3). At two timepoints during exponential growth, we obtained similar metabolite ^13^C-labeling patterns, which confirmed pseudo-steady state isotopic enrichment (Fig. 3). To elucidate fluxes through 48 reactions in the metabolic pathway, we combined the metabolite labeling data with substrate consumption rates (accounting for excretion of gluconate and 2-ketogluconate) and biomass growth rates (Fig. 4A; Fig. S2; Table S3; Table S4). Adjusting for the carbon loss through metabolite secretions, about 64% of the glucose consumed from the extracellular medium was available for intracellular catabolism in *P. protegens* Pf-5. Very close alignment between the MFA-estimated labeling patterns and those determined experimentally reflected the quality of the optimization procedure for the model cellular fluxes (Fig. S3).

**FIG 3.**
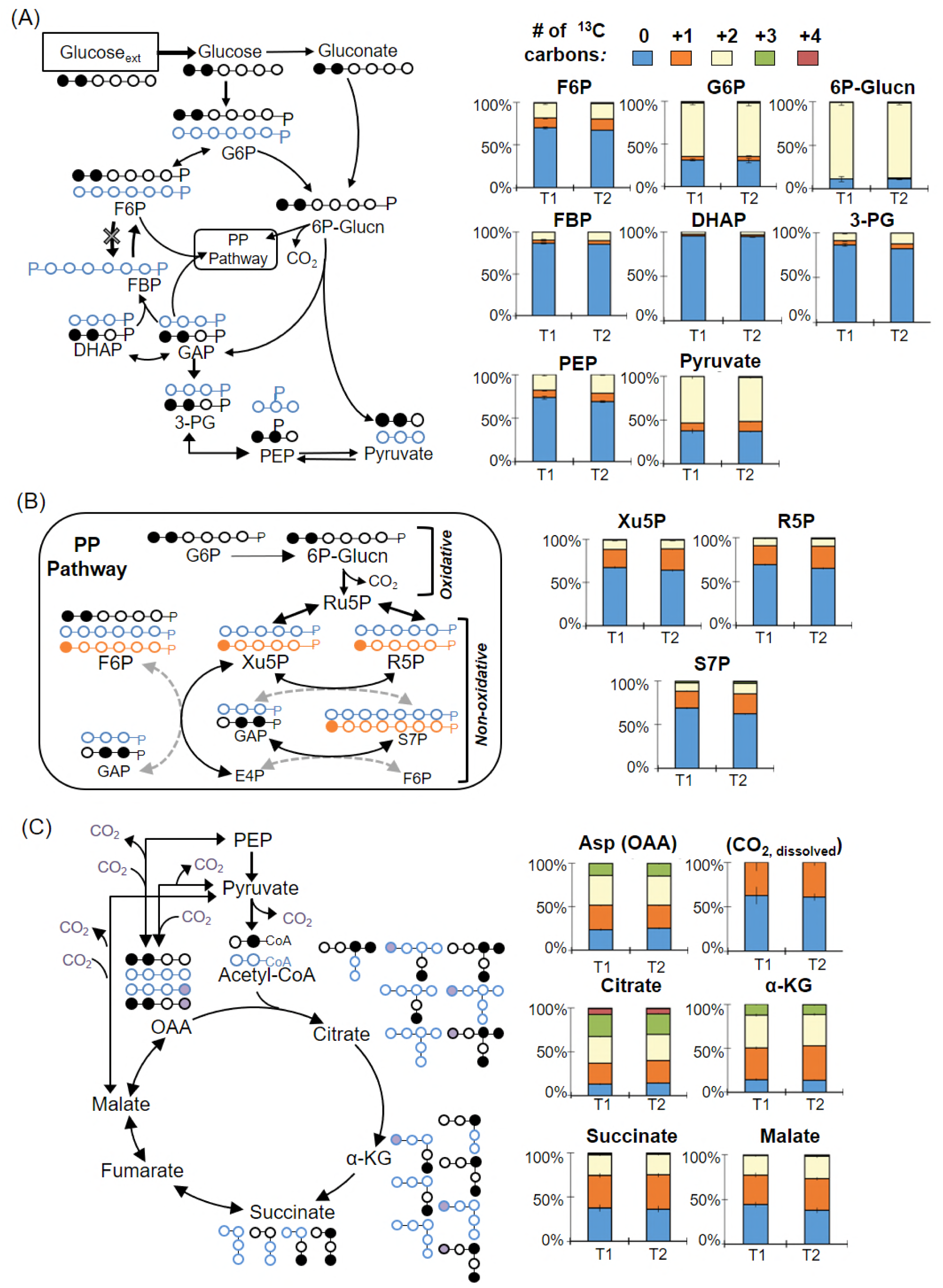
Long-term isotopic enrichment with [1,2-^13^C_2_]-glucose for carbon mapping of the metabolic network structure in *P. protegens* Pf-5. Carbon mapping is illustrated on the left and the metabolite labeling data are provided on the right for the following: (A) Initial glucose catabolism, Embden-Meyerhof-Parnas (EMP) pathway, and the Entner-Doudoroff (ED) pathway; (B) Oxidative and non-oxidative routes of the pentose-phosphate (PP) pathway; (C) the tricarboxylic acid (TCA) cycle. The dashed arrows describe minor formation routes of the metabolites. In the carbon mapping, blue represents the fate of structures derived from the ED and reverse-EMP pathways and orange represents the fate of structures derived from the oxidative PP pathway. Labeling patterns: nonlabeled (light blue), singly labeled (orange), doubly labeled, (cream), triply labeled (green), and quadruply labeled (red). Labeling data (average ± standard deviation) were from three independent biological replicates. The abbreviations are as follows: gluconate, Glucn; 2-ketogluconate, 2-KGlucn; glucose 6-phosphate, G6P; 6-phosphogluconate, 6P-Glucn; fructose 6-phosphate, F6P; fructose 1,6-bisphosphate, FBP; dihydroxyacetone-3-phosphate, DHAP; glyceraldehyde 3-phosphate, GAP; phosphoenolpyruvate, PEP; 3-phosphoglycerate, 3PG; xylulose 5-phosphate, Xu5P; ribose 5-phosphate, R5P; sedoheptulose 7-phosphate, S7P; oxaloacetate, OAA; aspartate, Asp; α-ketogluconate, α-KG.

**FIG 4.**
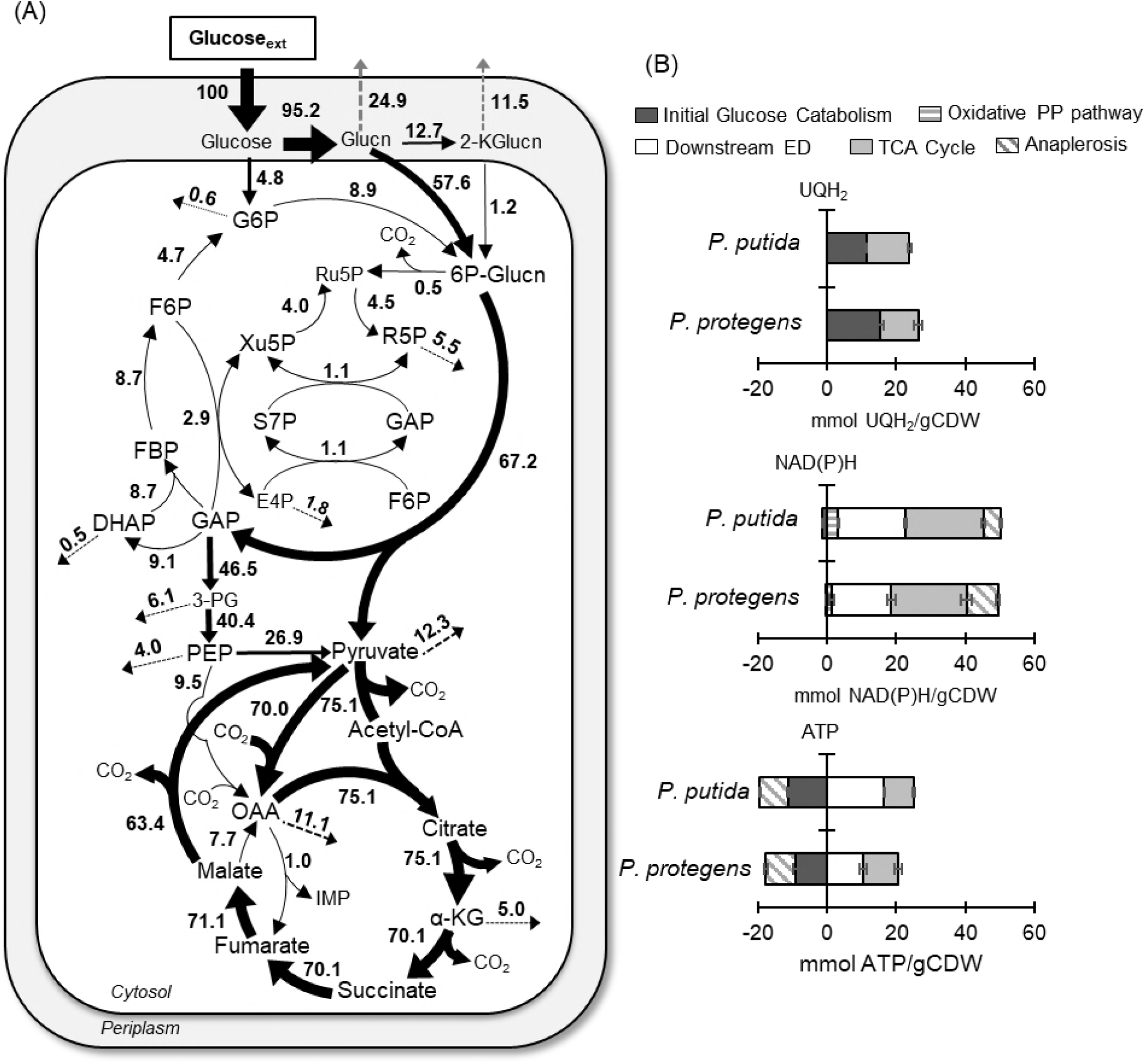
(A) Quantitative metabolic flux analysis and (B) energetic distributions in glucose-grown *P. protegens* Pf-5. All fluxes are normalized to 100% glucose uptake and the thickness of each arrow is scaled to the relative flux percentage. For A, black dotted arrows indicate contribution to biomass and grey dashed arrows indicate excretion fluxes. The absolute fluxes (mean ± standard deviation) are listed in Table S3 and S4. Abbreviations for A are as shown in the legend of Fig. 3. For B, the relative contribution of different pathways to the NAD(P)H, UQH_2_, and ATP pools were calculated from metabolic fluxes. Data for *P. putida* KT2440 were obtained from MFA reported in Nikel et al. (10). All values in part B are relative to glucose consumption rate and cellular biomass.

#### Involvement of glucose oxidation and the ED pathway

Glucose catabolism can be initiated via three different routes: direct phosphorylation to G6P, oxidation to gluconate in the periplasm before phosphorylation to 6P-gluconate, or additional periplasmic oxidation of gluconate to 2-ketogluconate before phosphorylation to 6P-gluconate (Fig. 1). Doubly ^13^C-labeled 6P-gluconate (86-89%) and doubly ^13^C-labeled G6P (∼63%) were both consistent with the assimilation of [1,2-^13^C_2_]-glucose (Fig. 3A). In accordance with the ED pathway wherein the first three carbons of 6P-gluconate become the doubly ^13^C-labeled pyruvate and the last three nonlabeled carbons become GAP, pyruvate was 50-53% doubly ^13^C-labeled and DHAP (an isomer of GAP) was over 95% nonlabeled (Fig. 3A). The 37-38% nonlabeled fraction of pyruvate was indicative of the nonlabeled fractions of precursor metabolites downstream of GAP (Fig. 3A). In the absence of the 6-phosphofructokinase enzyme in *P. protegens* Pf-5 and thus the lack of the traditional forward EMP pathway (7), the nonlabeled GAP and DHAP combined to produce the highly nonlabeled FBP (85-87%), which then cycled backward through upper EMP to result in the 67-70% nonlabeled F6P and 31% nonlabeled G6P (Fig. 3A). The MFA quantified the relative contribution of the three assimilation routes for the glucose carbons in *P. protegens* Pf-5 (Fig 4A). Only ∼5% of the glucose uptake was directly converted to G6P in the cytosol, whereas up to 95.2% of glucose was oxidized to gluconate accompanied by another flux (12.7%) toward 2-ketogluconate (Fig. 4A). Following the phosphorylation of the oxidized products of glucose to 6P-gluconate, our MFA determined that the flux through the ED pathway was 67.2% of the glucose uptake rate in *P. protegens* Pf-5 (Fig. 4A).

#### Oxidative versus non-oxidative PP pathway

Following a decarboxylation reaction through the oxidative PP pathway, doubly ^13^C-labeled 6P-gluconate would become singly ^13^C-labeled Ru5P, which would introduce singly ^13^C-labeled fractions into metabolites in the PP pathway (Fig. 3B). Thus, singly ^13^C-labeled fractions of xylulose 5-phosphate (Xu5P) (21-25%), ribose 5-phosphate (R5P) (22-25%), and sedoheptulose 7-phosphate (S7P) (20-23%) were due to flux through the oxidative PP pathway (Fig. 3B). However, by involving nonlabeled metabolites from downstream of the ED pathway, the non-oxidative PP pathway introduced relatively higher nonlabeled fractions of Xu5P (64-67%), R5P (65-69%), and S7P (62-69%) (Fig. 3B). Accordingly, the MFA determined an oxidative flux to the PP pathway (about 0.5%), which was a tenth of the non-oxidative PP pathway fluxes (5.1%) from ketolase and transaldolase reactions (Fig. 3B and 4A). As a precursor to both the oxidative PP pathway and the ED pathway, 6P-gluconate represents an important branch point in metabolism. Therefore, the low contribution of 6P-gluconate to the oxidative PP pathway necessitated a reverse flux from downstream ED pathway to the upper EMP pathway (i.e. gluconeogenesis) to channel glucose-derived carbons towards the PP pathway in support of biomass growth (Fig. 4A).

#### Downstream metabolic pathways

The ^13^C-labeling patterns of TCA cycle intermediates were consistent with the established route of carbon flow through this pathway (Fig. 3C). The decarboxylation of pyruvate generated nonlabeled and singly ^13^C-labeled acetyl moieties in acetyl-CoA, which were subsequently incorporated into the TCA cycle by combining with oxaloacetate (OAA) (nonlabeled, singly, doubly, and minorly triply ^13^C-labeled) to produce citrate (nonlabeled, singly, doubly, triply, and minorly quadruply ^13^C-labeled) (Fig 3C). The two sequential decarboxylation reactions in the TCA cycle led to the disappearance of the quadruply ^13^C-labeled fraction in citrate and, thereafter, the triply ^13^C-labeled fraction in α-ketoglutarate (Fig. 3C). The resulting succinate labeling pattern (nonlabeled, singly, and doubly ^13^C-labeled) led to a similar labeling scheme through fumarate, malate, and OAA (Fig. 3C). The MFA obtained a substantial flux (above 70% of the glucose uptake) through the TCA cycle from OAA around to malate (Fig. 4A).

Anaplerotic reactions contributed to the triply ^13^C-labeled OAA (Fig. 3C). The aforementioned decarboxylation reactions in the TCA cycle contributed to the ^13^C-labeled carbon dioxide (CO_2_) pool, which was calculated to be about 37-39% of the total dissolved CO_2_ (Fig. 3; Fig. S4). The carboxylation of doubly ^13^C-labeled pyruvate or doubly ^13^C-labeled PEP with ^13^C-labeled CO_2_ would generate triply ^13^C-labeled OAA (Fig. 3C). Notably, singly ^13^C-labeled OAA can be formed from carboxylation reactions of nonlabeled pyruvate or PEP with singly ^13^C-labeled CO_2_ (Fig. 3C). The singly ^13^C-labeled OAA can also be formed from singly ^13^C-labeled malate though the traditional TCA pathway utilizing malate dehydrogenase (Fig. 3C). The relative contributions of the different precursors to OAA (i.e., pyruvate/PEP versus malate) in *P. protegens* Pf-5 was resolved with the MFA, which demonstrated a substantially higher fractional flux of pyruvate to OAA (70%) than the flux of malate to OAA through malate dehydrogenase (7.7%) (Fig. 4A). This low flux through malate dehydrogenase was accompanied by a high flux for the direct conversion of malate to pyruvate (63.4%), thus highlighting a very active pyruvate shunt in *P. protegens* Pf-5 (Fig. 4A). The ^13^C-labeling patterns of TCA cycle metabolites implied an inactive glyoxylate shunt, which bypasses the decarboxylation reactions in the canonical TCA cycle to produce malate and succinate from citrate (Fig. S5). Specifically, there was a lack of triply ^13^C-labeled succinate which would be produced from triply and quadruply ^13^C-labeled citrate through the glyoxylate shunt (Fig. S5).

#### Energetics of glucose catabolism

We compared the energetic yields [reduced ubiquinone (UQH_2_), NAD(P)H, and ATP] from central carbon metabolism between our MFA-based cellular fluxes in *P. protegens* Pf-5 and those previously reported for *P. putida* KT2440, a well-studied biocatalyst candidate (10) (Fig. 4B). Compared to *P. putida* KT2440 (10), there was a slightly higher flux (about 5% higher) from glucose to gluconate in *P. protegens* Pf-5 but the flux from malate to OAA was lower (by about 25%) in *P. protegens* Pf-5. Accordingly, there was a higher yield of UQH_2_ from initial glucose catabolism in *P. protegens* Pf-5 but a higher UQH_2_ yield from the TCA cycle in *P. putida* KT2440 (Fig. 4B). Regarding the yield of NAD(P)H, there was a higher contribution from the TCA cycle (by about 3%) and from the oxidative PP pathway (by about 60%) in *P. putida* KT2440 than in *P. protegens* Pf-5 but the contribution of the anaplerotic reaction from malate to pyruvate was lower (by about 81%) in *P. putida* KT2440 than in *P. protegens* Pf-5 (Fig. 4B). With respect to ATP production by substrate-level phosphorylation, *P. protegens* Pf-5 produced less (by about 3 mmol ATP/g_CDW_) than *P. putida* KT2440 due to the relatively lower fluxes in the downstream ED pathway of *P. protegens* (10) (Fig. 4). However, the net ATP yield was similar because *P. protegens* Pf-5 consumed less ATP than *P. putida* KT2440 in initial glucose catabolism due to the higher flux for the glucose oxidation to gluconate and 2-ketogluconate and accounting for the subsequent carbon loss through secretions of these oxidized products in *P. protegens* Pf-5 (10) (Fig. 4). In sum, despite the different contributions of the relevant metabolic pathways, the combination of these contributions led to nearly equivalent net energetic yields in *P. protegens* Pf-5 and *P. putida* KT2440 (Fig. 4B).

### Hierarchy of Glucose Metabolism in the Presence of Other Carbohydrates

#### Proof of concept with ^13^C-labeled glucose and unlabeled glucose

Before determining the relative incorporation of [U-^13^C_6_]-glucose in the presence of a nonlabeled carbohydrate (xylose, arabinose, galactose, fructose, or mannose), we first conducted proof-of-concept experiments with the cells grown on [U-^13^C_6_]-glucose alone or in a 1:1 mixture with unlabeled glucose (Fig. 5; Fig. 6) (24). By comparative analysis with the metabolite labeling patterns in the latter two conditions, we seek to determine the relative incorporation of other unlabeled carbohydrates in the presence of [U-^13^C_6_]-glucose (Fig. 5; Fig. 6).

**FIG 5.**
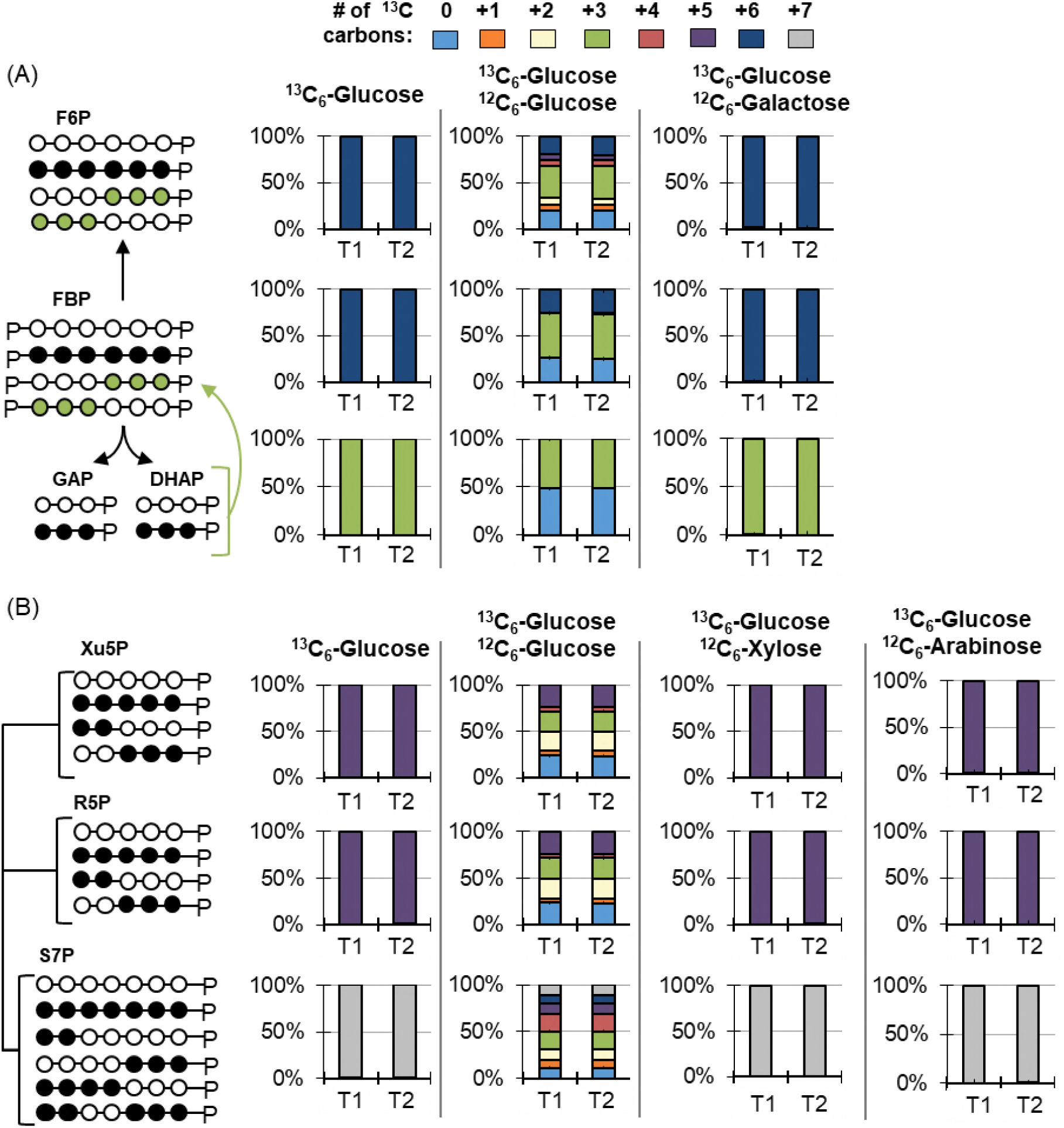
Metabolite labeling patterns during growth on [U-^13^C_6_]-glucose (^13^C_6_-Glucose) alone or with unlabeled glucose (^12^C_6_-Glucose), unlabeled galactose (^12^C_6_-Galactose), unlabeled xylose (^12^C_6_-Xylose), or unlabeled arabinose (^12^C_6_-Arabinose). (A) Carbon mapping (left) and the labeling data (right) for intracellular metabolites in the reverse Emden-Meyerhof-Parnas pathway following feeding on ^13^C_6_-Gluc alone or with ^12^C_6_-Gluc or ^12^C_6_-Gala. (B) Carbon mapping (left) and the labeling data (right) for intracellular metabolites in the pentose-phosphate pathway following feeding on ^13^C_6_-Gluc alone or with ^12^C_6_-Gluc, ^12^C_6_-Xylo, or ^12^C_6_-Arab. In the carbon mapping, the open circles and the filled circles represent unlabeled and ^13^C-carbons, respectively. Data were obtained at two timepoints during exponential growth: at OD_600_ 0.5-0.6 (T1) and at OD_600_ 0.9-1.0 (T2). Labeling color legend: nonlabeled carbon (light blue), one ^13^C-carbon (orange), two ^13^C-carbons (cream), three ^13^C-carbons (green), four ^13^C-carbons (red), five ^13^C-carbons (purple), six ^13^C-carbons (dark blue), and seven ^13^C-carbons (grey). Labeling data (average ± standard deviation) were from three biological replicates. Very small error bars are not noticeable. The abbreviations used for the metabolites are given in Fig. 1 and Fig. 3.

**FIG 6.**
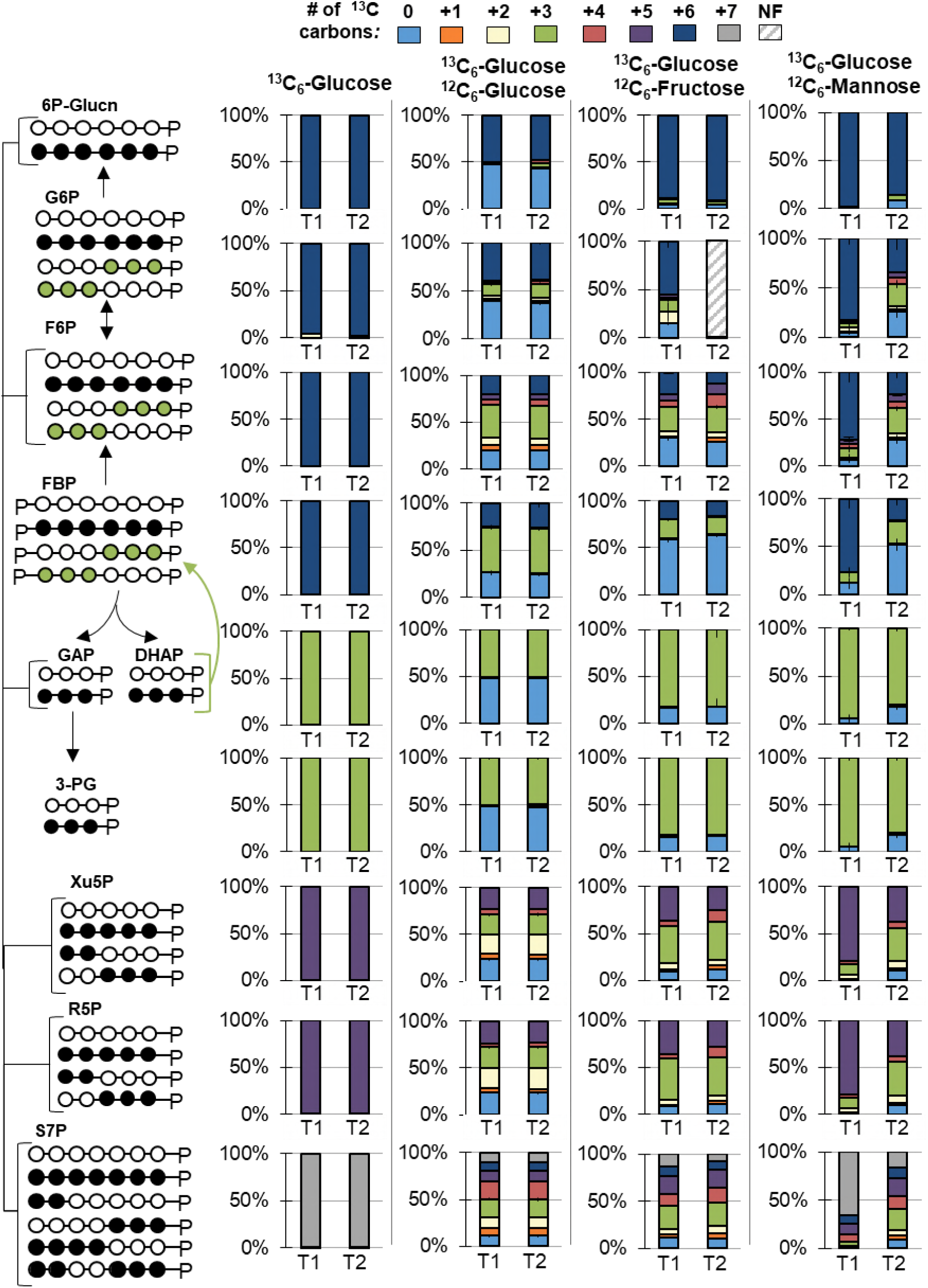
Metabolite labeling patterns during growth on [U-^13^C_6_]-glucose (^13^C_6_-Glucose) alone or with unlabeled glucose (^12^C_6_-Glucose), unlabeled fructose (^12^C_6_-Fructose), or unlabeled mannose (^12^C_6_-Mannose). Carbon mapping (left) and the labeling data (right) for intracellular metabolites in the Entner-Doudoroff pathway, reverse Emden-Meyerhof-Parnas (EMP) pathway, and the pentose-phosphate pathway. The open circles and the filled circles represent unlabeled and ^13^C-carbons, respectively; the-green colored circles represent labeling schemes specifically from reverse flux through upper EMP pathway. Data were obtained at two timepoints during exponential growth: at OD_600_ of 0.5-0.6 (T1) and at OD_600_ of 0.9-1.0 (T2). Labeling color legend: nonlabeled carbon (light blue), one ^13^C-carbon (orange), two ^13^C-carbons (cream), three ^13^C-carbons (green), four ^13^C-carbons (red), five ^13^C-carbons (purple), six ^13^C-carbons (dark blue), and seven ^13^C-carbons (grey). NF, not found. Labeling data (average ± standard deviation) were from biological replicates (*n* = 3). Very small error bars are not noticeable. The abbreviations used are given in the legend of Fig. 1 and Fig. 3.

#### Glucose with xylose, arabinose, or galactose

While galactose, xylose, and arabinose are reported to not be metabolized as single substrates in *P. protegens* Pf-5 (6), whether they are metabolized in the presence of glucose needed to be evaluated. During growth on ^13^C-labeled glucose and galactose, the labeling patterns of metabolites in the EMP and ED pathways (F6P, FBP, DHAP) were identical to the metabolite labeling during feeding on glucose alone, thus indicating the lack of galactose catabolism in the presence of glucose (Fig. 5A). During simultaneous feeding on ^13^C-labeled glucose and a pentose substrate (xylose or arabinose), the labeling patterns of metabolites in the PP pathway (Xu5P, R5P, S7P) also indicated the lack of pentose assimilation (Fig. 5B). Furthermore, there was no indication of xylose incorporation into α-ketoglutarate, consistent with the lack of the Weimberg pathway (Fig. S6). Therefore, our data from ^13^C enrichment studies ascertained the absence of carbon assimilation from galactose, xylose, or arabinose in the presence of glucose (Fig. 5).

#### Glucose with Fructose

Following simultaneous feeding on [U-^13^C_6_]-glucose and unlabeled fructose, a persistence of nonlabeled fractions in the intracellular metabolites was consistent with both uptake and assimilation of fructose in the presence of glucose (Fig. 6). However, the differential abundance of the nonlabeled fractions indicated a bottleneck in fructose assimilation, which may explain the slower rate of fructose depletion than glucose depletion from the extracellular medium (Fig. 2B; Fig. 6). Consistent with fructose incorporation through F1P into FBP, the highest fraction of nonlabeled carbons was seen in FBP (59-62%); both F6P and DHAP had lower nonlabeled fractions (25-30% and 17%, respectively) (Fig. 6). The lower fraction of nonlabeled carbons in DHAP than in F6P implied that the fructose-derived carbons were preferentially routed through a backward flux through upper EMP pathway towards the ED pathway (Fig. 6). The labeling of G6P reflected the nonlabeled and partially ^13^C-labeled fractions from F6P, consistent with this backward flux (Fig. 6). Therefore, in lieu of the forward EMP pathway, our data stresses the importance of the ED pathway in the co-processing of fructose with glucose.

The labeling patterns of metabolites in the PP pathway also showed that the contribution of the fructose-derived carbons in this pathway was preferentially through the non-oxidative route (Fig. 6). A transketolase reaction in the non-oxidative PP pathway combines the first two carbons of F6P with GAP to produce Xu5P. Both Xu5P and R5P have significant fractions of triply ^13^C-labeled carbons (38-44%), in accordance with the combination of nonlabeled F6P with triply ^13^C-labeled GAP following growth on ^13^C-labeled glucose with unlabeled fructose (Fig. 6). Due to the low fraction of nonlabeled GAP (as determined from DHAP labeling), there was a lack of appreciable fraction of doubly ^13^C-labeled R5P and Xu5P, which was evident in cells grown on [U-^13^C_6_]-glucose with unlabeled glucose (Fig. 6).

#### Glucose with mannose

In contrast to the metabolite labeling during growth on [U-^13^C_6_]-glucose with unlabeled fructose, the metabolite labeling patterns following feeding on the [U-^13^C_6_]-glucose with unlabeled mannose showed delayed assimilation of mannose into intracellular metabolism during exponential cell growth (Fig. 6). This was evident by the substantial increase in the nonlabeled fraction of FBP at the two timepoints, from 12% at an optical density at 600 nm (OD_600_) of ∼0.5 to 52% at OD_600_ of ∼1.0 (Fig. 6). This time-dependent labeling data agreed with the fact that extracellular mannose started to decrease significantly after 6 h of growth, following the depletion of glucose (Fig. 2B).

A higher nonlabeled fraction of FBP than of F6P implied that incorporation of mannose-derived carbons into FBP by way of fructose was preferred in *P. protegens* Pf-5 (Fig. 6). In agreement with this catabolic route for mannose, there was a larger pool of FBP than F6P when cells were grown on the glucose:mannose mixture relative to the glucose only condition (Fig. S7). This large FBP pool was also seen in the glucose:fructose condition relative to the glucose only condition (Fig. S7). During feeding on ^13^C-glucose with unlabeled mannose, the nonlabeled fraction of DHAP (18%) was lower than F6P (28%) and G6P (26%) (Fig. 6). Therefore, these labeling data collectively indicated the cycling of nonlabeled mannose carbons backward through the upper EMP pathway towards the ED pathway, similar to the intracellular metabolism of fructose (Fig. 6).

### Cellular Carbon Fluxes during Co-utilization of Glucose and Fructose

Quantitative MFA was used to assess differences in the metabolic fluxes during growth on glucose alone versus growth on glucose with fructose (Fig. 7; Table S4; Table S5). Our MFA focused on the pathways surrounding the incorporation points of carbohydrates: initial glucose catabolism, the EMP pathway, PP pathway, and the ED pathway (Fig. 7). Each MFA made use of the labeling data collected at OD_600_ of 0.5 during early exponential phase before extracellular glucose is depleted and was constrained by the consumption rate of each carbohydrate, biomass effluxes, and metabolite secretions (Fig. 7; Fig. S2; Table S5; Table S6). Upon optimization of the estimated metabolic fluxes, the model-estimated ^13^C-labeling patterns agreed well with the experimentally determined ^13^C-labeling patterns for each condition (Fig. S8; Fig. S9).

**FIG 7.**
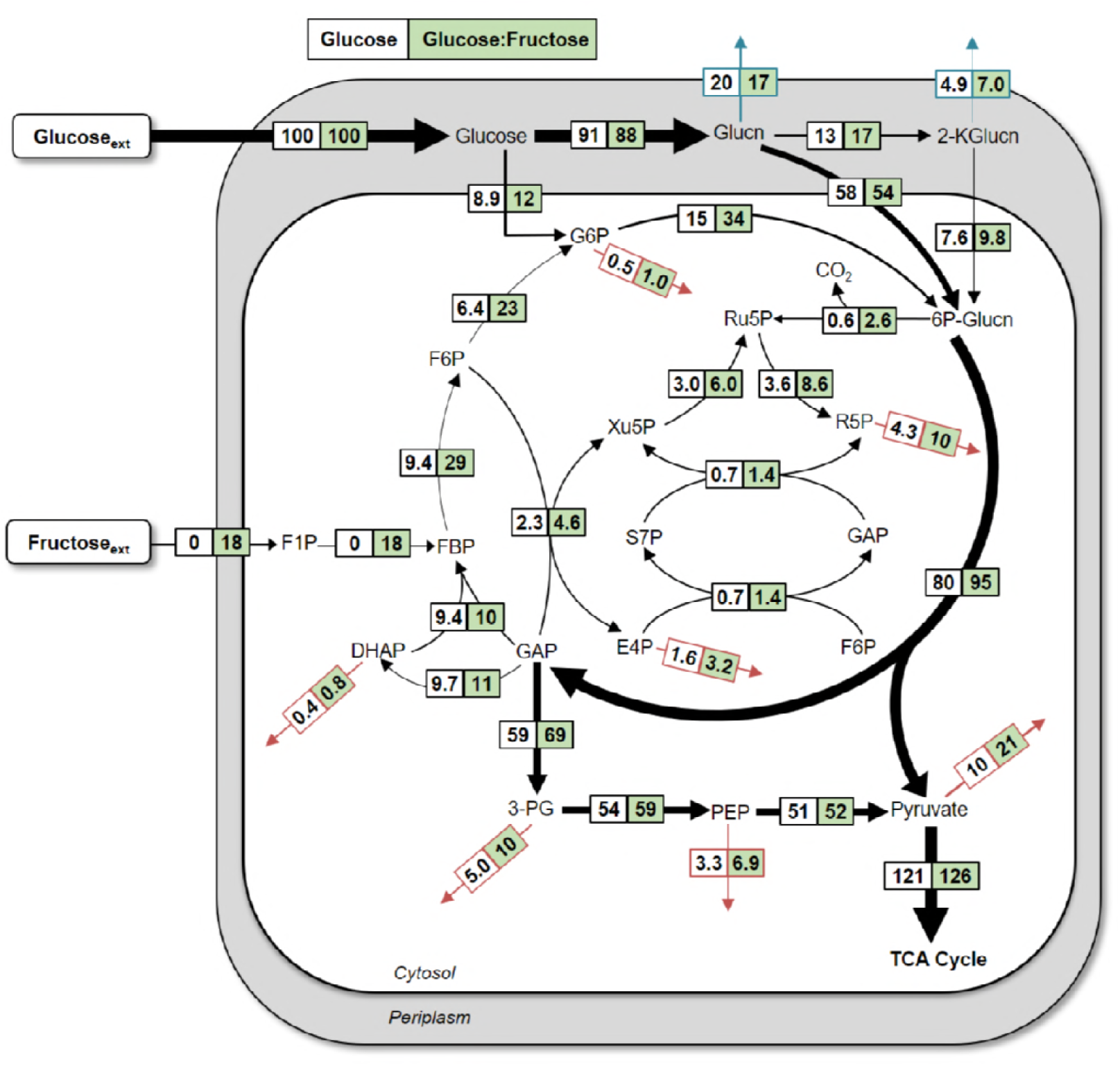
Quantitative metabolic flux analysis of *P. protegens* Pf-5 using metabolite labeling data following growth on [U-^13^C_6_]-glucose:unlabeled glucose (white) or [U-^13^C_6_]-glucose:unlabeled fructose (green). All fluxes were normalized to 100% glucose uptake and the thickness of each arrow was scaled to the relative flux percentage for the glucose-only growth condition. The absolute fluxes (mean ± standard deviation) are listed in Table S5 and S7. Red arrows indicate contribution to biomass and blue arrows indicate excretion fluxes. The metabolite abbreviations are as given in the legends of Fig. 1 and Fig. 3.

Despite the genome-encoded capabilities in *P. protegens* Pf-5 to involve the splitting of FBP to DHAP and GAP (Paulsen et al., 2005), the MFA revealed that the net flux was instead the aldolase reaction that combines DHAP and GAP to generate FBP (Fig. 7). Due to the additional incorporation of fructose-derived carbons through the flux from FBP to 6P-gluconate, there were higher fluxes from FBP to F6P (3-fold increase), F6P to G6P (3.6-fold increase), and G6P to 6P-gluconate (2.3-fold increase) in cells grown on the glucose:fructose mixture compared to glucose alone (Fig. 7). Similar to metabolism of glucose alone, there was a significant flux of the glucose uptake channeled through glucose oxidation to gluconate (88%) followed by the ED pathway (95%) during the metabolism of both glucose and fructose (Fig. 7). Consistent with the increased carbon flux towards 6P-gluconate, there were a 4.3-fold increase in the flux toward the oxidative PP pathway (i.e. from 6P-gluconate to Ru5P) and a 19% increase in the ED pathway (i.e., from 6-gluconate to GAP and pyruvate) (Fig. 7).

### Kinetic Isotope Analysis during Co-utilization of Glucose and Mannose

Due to the lack of isotopic pseudo-steady state during simultaneous growth on glucose and mannose, ^13^C-MFA could not be conducted. To capture the assimilation route for mannose, we obtained measurements of metabolic isotopic fractions during six timepoints during exponential growth of ^13^C-labeled glucose and unlabeled mannose (Fig. 8). Specifically, we examined the labeling patterns of gluconate, 6P-gluconate, G6P, F6P, FBP, and DHAP (Fig. 8). Across all timepoints, gluconate labeling was consistently ∼100% fully ^13^C-labeled, indicating that gluconate was exclusively the oxidized product of the ^13^C-labeled glucose (Fig 8). However, the labeling of the other five metabolites had nonlabeled fractions derived from the assimilation of unlabeled mannose (Fig. 8). The labeling of 6P-gluconate exhibited the slowest kinetic incorporation of nonlabeled fractions (Fig. 8). The appearance of nonlabeled pool of 6P-gluconate (starting at ∼4%) occurred at the fifth measurement timepoint at an OD_600_ of 0.94 (Fig. 8). By contrast, FBP exhibited the fastest incorporation of nonlabeled fraction that occurred at an OD_600_ of 0.2 (Fig. 8). Compared to the labeling kinetics of FBP, there was a delay in the incorporation of nonlabeled carbons in F6P and G6P, which started to occur an OD_600_ of 0.6, and DHAP, which steadily increased after an OD_600_ 0f 0.8 (Fig. 8). Statistical analysis (Mixed Effect Model; F_3, 57_ = 12.973; p < 0 .0001) confirmed that the significant effect of OD_600_ on the incorporation of mannose-derived nonlabeled carbons was dependent on the metabolite (Fig. 8). These kinetics data collectively demonstrated that, instead of being channeled directly from FBP to GAP and DHAP, mannose carbons were incorporated at FBP and cycled up through F6P and G6P to the ED pathway to generate subsequently GAP and DHAP (Fig. 8). Thus, the catabolic route for mannose was similar to what was determined for fructose catabolism.

**FIG 8.**
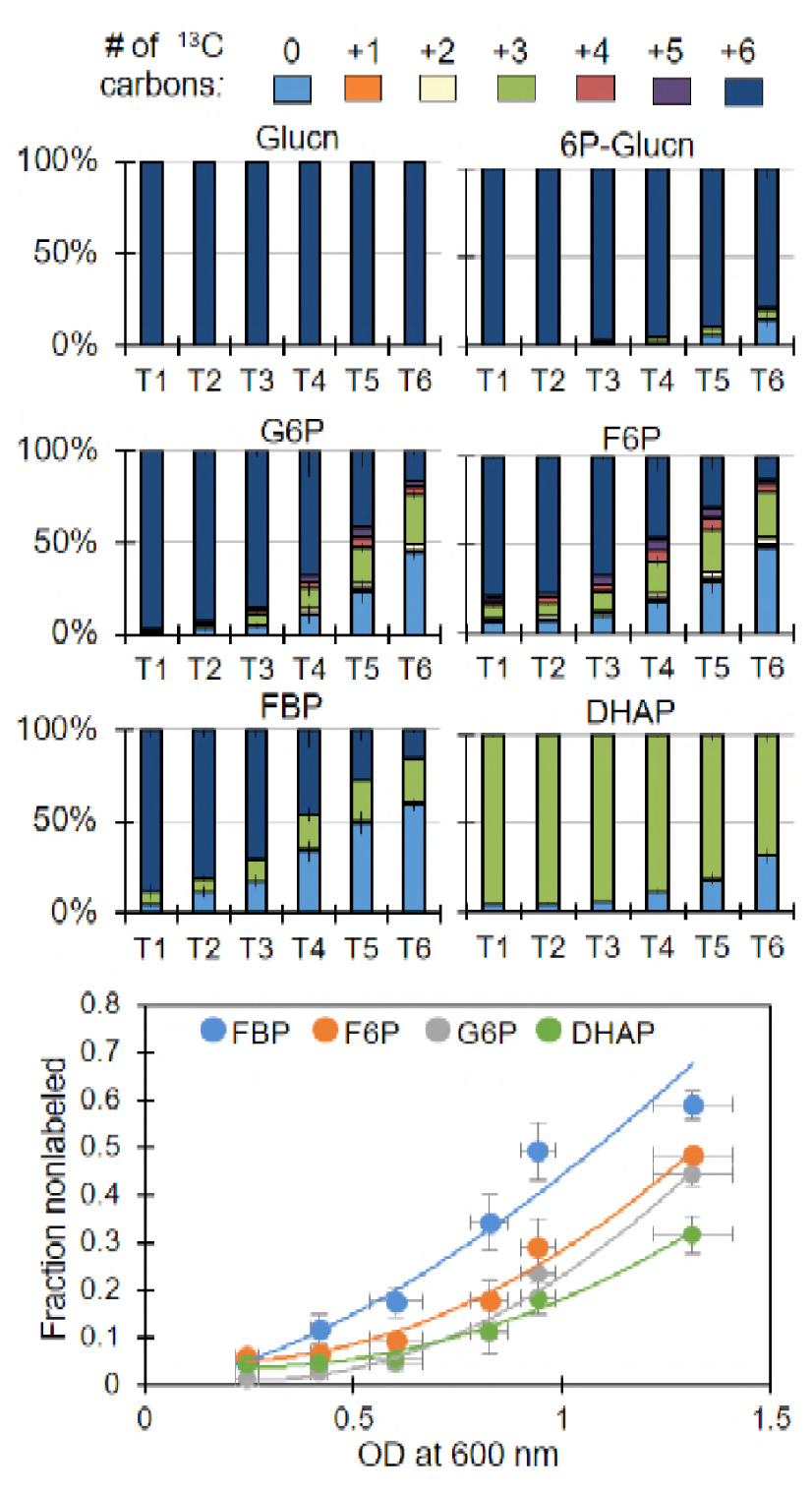
Kinetic profiling of metabolite labeling patterns in cells grown on [U-^13^C_6_]-glucose and unlabeled mannose. Labeling color legend: nonlabeled carbon (light blue), one ^13^C-carbon (orange), two ^13^C-carbons (cream), three ^13^C-carbons (green), four ^13^C-carbons (red), five ^13^C-carbons (purple), six ^13^C-carbons (dark blue). Data were obtained at six timepoints during exponential growth: at OD_600_ of 0.21-0.27 (T1), at OD_600_ of 0.40-.44 (T2), at OD_600_ of 0.54-0.67 (T3), at OD_600_ of 0.79-.87 (T4), at OD_600_ of 0.90-0.98 (T5), and at OD_600_ of 1.2-1.4 (T6). The data (average ± standard deviation) are from biological replicates (*n* = 3). The lines through the data in the bottom figure represent guide to the eye. The metabolite abbreviations are as given in the legends of Fig. 1.

## DISCUSSION

The metabolic networks and fluxes of several *Pseudomonas* species, including *P. putida, P. fluorescens,* and *P. aeruginosa*, have been previously described (8–11, 22, 25). Here we present the first metabolic flux analysis of the recently characterized *P. protegens* Pf-5, a common plant commensal bacterium known to secrete specialized metabolites important for the biocontrol of fungi and bacteria pathogenic to plants (4, 6). Metabolic flux quantitation determined that initial glucose catabolism in *P. protegens* Pf-5 was primarily through periplasmic oxidation to gluconate with relatively minor influx of glucose through G6P (Fig. 4A). Up to 95% of consumed glucose in *P. protegens* Pf-5 was channeled through the ED pathway, which was also reported in *P. putida, P. fluorescens,* and *P. aeruginosa* (8, 11, 22, 25). Furthermore, the non-oxidative route was more significant than the oxidative route in generating PP pathway intermediates in *P. protegens* Pf-5, as previously reported for *P. putida* KT2440 (11) (Fig. 3B; Fig. 4A). The highly active flux of pyruvate formation from malate in the *P. protegens* Pf-5 cells was also reported in *P. putida* and *P. fluorescens* (8, 10, 22). In sum, our results stressed that the metabolic network for glucose metabolism in *P. protegens* Pf-5 is consistent with the metabolic network of previously studied species of the *Pseudomonas* genus. Finally, through the contribution of different metabolic pathways, the total energetic yields of *P. protegens* Pf-5 were remarkably similar to *P. putida* KT2440, whose metabolism was been featured for its capability for fulfilling high demands of reducing power (10).

Root exudates and the breakdown of polysaccharides from plant biomass both provide various carbohydrates that stimulate growth of soil microorganisms, including *Pseudomonas* species (26). With respect to the catabolism of other carbohydrates besides glucose, we found that *P. protegens* Pf-5 did not metabolize the common hemicellulose monomers galactose, xylose, or arabinose in the presence of glucose but did utilize the carbon mixtures of glucose with fructose or mannose (Fig. 5; Fig. 6). In accordance with gene annotations, metabolite labeling data confirmed that fructose was incorporated into metabolism via FBP (Fig. 6). However, contrary to the possible route for mannose assimilation through F6P annotated in the *P. protegens* Pf-5 genome (7), the primary route of mannose assimilation in *P. protegens* Pf-5 was found to be also via FBP (Fig. 6). In addition, the appearance of fructose extracellularly during growth on mannose implied conversion of mannose to fructose prior to intracellular metabolism (Fig. 2B). Mannose conversion to fructose by a mannose isomerase has been reported previously in *P. cepacia, P. aeruginosa,* and *P. saccharophila* (17–20). Whether a non-specific isomerase exists in *P. protegens* remains to be determined.

The cyclic metabolism linking backward flux from FBP through the upper EMP towards the ED pathway produces equivalent NADPH, but less net ATP, compared to direct FBP conversion to GAP and DHAP. Interestingly, instead of this direct contribution to the lower EMP pathway, carbons from assimilated fructose and mannose were routed through the cyclic connection between upper EMP pathway and the ED pathway during mixed-substrate utilization (Fig. 7; Fig. 8). Comparable MFA findings were reported for fructose-only catabolism in *P. putida* KT2440 and *P. fluorescens* SBW25 (14, 16). Therefore, *P. protegens* Pf-5 exhibits a strong reliance on the ED pathway for both fructose and mannose assimilation even in the presence of glucose.

Similar growth phenotypes during co-utilization of hexoses implied that, despite different carbohydrates in the growth medium at the same total carbon equivalence, *P. protegens* Pf-5 preserved a constant biomass maintenance (Fig. 2A). However, uptake of glucose was preferred over the uptake of fructose (3 to 1) or mannose (4 to 1) (Fig. 2; Table S2). The composition ratio of glucose to fructose in maize root exudates was found to be 2 to 1 (27). And, across soil horizons, the glucose:mannose ratio ranged approximately from 3:1 to 5:1 (28). Therefore, remarkably, the relative consumption rates of glucose versus fructose or mannose in *P. protegens* Pf-5 were in agreement with relative composition of these carbohydrates in environmentally-relevant conditions.

Related to the potential of *P. protegens* Pf-5 as a biocatalytic platform, the innate production and secretion of gluconate and 2-ketogluconate in *P. protegens* Pf-5 are additional attractive features. Oxidized sugars are important precursors to polymeric materials including polyesters. In fact, gluconate was identified as a top 30 value-added candidate for production of bio-inspired materials (29). Under our experimental conditions, the secretion rates of gluconate and 2-ketogluconate collectively accounted for about 35% of the glucose uptake in *P. protegens* during exponential growth whereas metabolite secretion of gluconate was reported to be less 10% of the glucose uptake in *P. putida* KT2440 (11) (Fig. 2C). While there were relatively similar secretions during growth on glucose alone or glucose and fructose, growth on glucose and mannose resulted in a decrease in the total secretion (by about 3.5 mM) of gluconate and 2-ketogluconate (Fig. 2C). After glucose was depleted, a decrease in the concentration of both gluconate and 2-ketogluconate indicated that the cells can utilize these metabolites once their favored carbon source is exhausted (Fig. 2B; Fig. 2C). This phenomenon would need to be considered and manipulated to harvest these metabolite secretions in *P. protegens* Pf-5 as valuable products. In light of the abundance of different types of carbohydrates in renewable carbon feedstocks in natural environments and in use for engineered bioproduction (30), our findings collectively provide important insights regarding the cellular metabolism underlying carbohydrate co-utilization in *P. protegens* Pf-5 and related *Pseudomonas* species.

## MATERIALS AND METHODS

### Materials

The *P. protegens* Pf-5 cells were acquired from the American Type Culture Collection (Manassas, VA). Unless noted otherwise, chemicals used in the growth media were obtained from Sigma-Aldrich (St. Louis, MO), Cayman Chemical (Ann Arbor, MI), or Fisher Scientific (Pittsburgh, PA). The ^13^C-labeled glucose ([U-^13^-C_6_]-glucose and [1,2-^13^-C_6_]-glucose) were purchased from Cambridge Isotopes (Tewskbury, MA) and Omicron Biochemicals (South Bend, IN), respectively. All culture solutions were prepared with Millipore water (18.2 MΩ cm, Millipore; Billerica, MA, USA) while resuspensions for LC-HRMS analysis were made with LC-MS grade water. Solutions were sterilized by passing through a 0.22-μm nylon filters (Waters Corporation, MA). An Agilent Cary UV-visible spectrophotometer (Santa Clara, CA) was used for optical density readings at 600 nm. The LC-HRMS analysis was conducted on an ultra-high-performance LC (Thermo scientific DionexUltiMate 3000) coupled to a high-resolution/accurate-mass MS (Thermo Scientific Q Exactive quadrupole-Oribitrap hybrid MS) with electrospray ionization.

### Culturing conditions and growth measurements

Batch growth experiments (three to seven biological replicates) of *P. protegens* Pf-5 were conducted in an incubator (model I24; New Brunswick Scientific, Edison, NJ) maintained at 30°C and shaken at 220 rpm. Initial growth in nutrient-rich medium prior to growth in minimal-nutrient medium was conducted as previously described (11). Final growth experiments were conducted in 125-mL baffled flasks with a pH-adjusted (7.0) and filter-sterilized minimal-nutrient medium that contained major salts and essential trace metal nutrients as previously reported (31): 89.4 mM K_2_HPO_4_, 56.4 mM NaH_2_PO_4_, 0.81 mM MgSO_4_·7H_2_O, 18.7 mM NH_4_Cl, 8.6 mM NaCl, 34 μM CaCl_2_·2 H_2_O, 30 μM FeSO_4_·7 H_2_O, 0.86 μM CuSO_4_·5 H_2_O, 1.9 μM H_3_BO_3_, 7.7 μM ZnSO_4_·7 H_2_O, 0.75 μM MnSO_4_ ·5 H_2_O, 0.26 μM NiCl_2_·6 H_2_O, and 0.3 1μM Na_2_MoO_4_·5 H_2_O. The carbohydrate composition was 100 mM C total for glucose alone (equivalent to 16.7 mM or 3 g L^-1^ glucose) and for 1:1 glucose:xylose, glucose:arabinose, glucose:galactose, glucose:mannose, or glucose:fructose. For cellular isotopic enrichment, either [U-^13^C_6_]-glucose or [1,2-^13^C_6_]-glucose was used in glucose-only growth, but only [U-^13^C_6_]-glucose was used for mixtures in combination with an unlabeled second carbohydrate. Bacterial growth in the biological replicates was monitored as a function of time until late stationary phase using OD_600_ measurements (Fig. S1)—cell suspensions were diluted when the OD_600_ value was above 0.5 to get accurate reading. Cell dry weight in grams (g_CDW_) was also determined throughout growth by lyophilizing the cell pellets as previously described (11).

### Measurement of Carbohydrate Consumption

Independent ^13^C-tracer experiments, as described in the next section, confirmed that extracellular depletion of the substrates correlated with substrate consumption. The extracellular depletion of each carbohydrate substrate (three biological replicates) was determined throughout 24 h of cell growth. Culture aliquots were pelleted by centrifugation and the supernatant was stored at −20 °C until further analysis. Following previously reported LC methods for carbohydrate analysis (21), we applied an analytical method using LC-HRMS for monitoring carbohydrate concentration in the extracellular solution. Peak identification and quantification of carbohydrate concentrations were conducted with ThermoScientific Xcalibur™ 3.0 Quan Browser.

### Metabolite Monitoring and Quantification

#### Extracellular Metabolites

To determine metabolite excretion rates, cell suspension samples (three biological replicates) were harvested periodically throughout growth and pelleted with centrifugation before the supernatant was removed and stored at −20 °C until LC-HRMS analysis. Dilutions of 1:10, 1:100, and 1:1000 were conducted to account for the varying concentrations of each metabolite over time. For the LC, an Acquity UPLC Waters 1.7 µm particle size column with dimensions of 2.1 x 100 mm was used for all metabolomics samples (Milford, MA) with constant column temperature of 25°C. The flow rate was kept constant at 0.180-mL min^-1^. The mobile phase composition and LC protocol were as previously described (31). The injection volume for each sample was 10-µL. The MS was operated in full scan negative mode. Metabolite identification was based on accurate mass and matches with standard retention time. Metabolite levels were quantified using ThermoScientific Xcalibur™ 3.0 Quan Browser.

#### Intracellular Metabolites

Cells were separated by filtration and then lysed to extract intracellular metabolites as described in Sasnow et al. (11). Metabolites in solution were monitored by LC-HRMS and the ^13^C labeling patterns were analyzed on the Metabolomic Analysis and Visualization Engine (MAVEN) software (32, 33). Isotopologue data were obtained for the following compounds: 6P-gluconate, G6P, F6P, FBP, DHAP, 3-phosphoglycerate, PEP, pyruvate, Xu5P, R5P, S7P, aspartate, citrate, α-ketoglutarate, succinate, and malate. Aspartate ^13^C-labeling is used as a proxy for OAA ^13^C-labeling by assuming equilibrium between the two compounds (11). The labeling of dissolved CO_2_ was estimated from the labeling patterns of ornithine and citrulline (Fig. S2); ornithine incorporates one mole of dissolved CO_2_ to become citrulline. All the extracted isotopologues were corrected for natural abundance of ^13^C. To verify pseudo-steady state isotopic enrichment of the intracellular pools, metabolites were isolated from cellular extracts obtained at two different timepoints during the exponential phase, at OD_600_ values of ∼0.5 and ∼1.0 (11). To analyze mannose incorporation over time, cells were extracted at six timepoints during exponential growth corresponding to OD_600_ values of ∼0.2, ∼0.4, ∼0.6, ∼0.8, ∼0.9, and ∼1.3. A mixed effect model was conducted using R (34) and the lmerTest package (35), which modeled the nonlabeled fraction (log transformed) by OD_600_, metabolite, and their interaction with the random effect of biological replicate.

### Quantitative Metabolic Flux Modeling

Quantitation of the metabolic fluxes was achieved for cells grown on glucose alone and glucose with fructose. We employed the following experimental data to constrain the metabolic flux analysis: substrate uptake rate, metabolite excretions, growth rate, and cellular stoichiometry. Carbon effluxes from intermediates in central metabolism towards biomass production were determined based on the growth rate for each condition and the biomass composition of *P. putida* (nucleic acids, proteins, cell membrane, and carbohydrate polymers) (36). An initial reaction network for the central carbon metabolism of *P. protegens* PF-5 was constructed using predicted genome-scale metabolic model (7), and gene annotation of metabolic enzymes reported on the KEGG database (37–39), and MetaCyc (40). The metabolic reaction network was validated through ^13^C-labeling of intracellular metabolites. The following reactions were constrained in the forward direction: gluconate → 6P-gluconate, 6P-gluconate → ribulose 5-phosphate, gluconate → 2-ketogluconate, 2-ketogluconate → 6P-gluconate, glucose → G6P, FBP → F6P, malate → pyruvate, pyruvate → OAA, and PEP → OAA. Optimized fluxes in the model metabolic network reactions were determined by the 13CFLUX2 software package (http://www.13cflux.net) (41) whereby quality of fit was optimized iteratively by comparing experimental ^13^C-labeling data and the *in silico*-estimated labeling data.

## SUPPLEMENTAL MATERIAL

Supplemental material can be found in the online version of this article.

## ACKNOWLEDGEMENTS

Graduate financial support for R.A.W and C.M.M. was provided in part by the College of Agriculture and Life Sciences at Cornell University. We are grateful to Dr. Krista A. Barzen-Hanson from the Aristilde Research Group at Cornell University (presently at Elmira College) for her technical assistance in implementing the LC-MS protocol to analyze the carbohydrate compounds in solution. We acknowledge Erika Mudrak at the Cornell Statistical Consulting Unit for her assistance with implementing and interpreting the statistical analysis.

